# Reproducible generation of Nipah virus pseudovirions with uniform expression of F and G surface glycoproteins for high-throughput neutralization assays

**DOI:** 10.1101/2025.04.07.647518

**Authors:** Geetu Rose Varghese, Vivek Vijay, Sreeja Sreedevi, Santhik S Lupitha, Priya Prabhakaran, Sushama Aswathyraj, Anitha P. Moorkoth, Niyas K Pulloor, Easwaran Sreekumar

## Abstract

Nipah virus (NiV) is a pathogen to be handled in BSL-4 facilities. Multiple surrogate systems such as virus-like particles (VLPs) and pseudoviruses enable carrying out NiV neutralization assays and study of virus entry pathways in biosafety level-2 (BSL-2) facilities. These are dual protein expression and surface display systems comprising NiV structural glycoproteins F and G as the key components. During generation of NiV VLPs or pseudovirions, ensuring batch to batch uniformity is a major concern due to the lack of proportionate expression of these proteins in the producer cells as well as their consistent incorporation in the particles. We established HEK293 pseudovirion producer cells that stably co-express NiV F and G proteins to address the issue. Fluorescence-activated cell sorting (FACS) analysis of clonally selected cells for high and uniform level F and G protein co-expression; and their further expansion were carried out to further refine the system. High titer vesicular stomatitis virus (VSV)-based pseudoviruses which showed consistent expression of both the glycoprotein were reproducibly generated from these producer cells. In functional assays, these pseudovirions exhibited a dose-dependent neutralization by commercial anti-NiV F and G antibodies as well as by convalescent serum from Nipah recovered patients. A pseudovirus neutralization test (PVNT) with a secreted alkaline phosphatase (SEAP) as the reporter was established in the study. The assay supports high-throughput adaptability with a quick turn-around time. It will aid large-scale human and animal serosurveillance studies in Nipah endemic regions as well as screening of virus entry inhibitors and monoclonal antibodies.

## Introduction

Nipah virus (NiV) is a highly lethal zoonotic paramyxovirus causing fatal encephalitis in humans [1]. Fruit bats of the *Pteropus* genus serve as a natural reservoir of the virus [2]. The initial Nipah cases identified in 1998 in Malaysia and Singapore were caused by spill over infections from pigs to humans [3–5]. On the other hand, a human-to-human transmission was evident in the re-emerging outbreaks of the disease in Bangladesh and neighbouring regions in India [6, 7]. In southern peninsular India the first outbreak occurred in 2018 in Kerala with 18 laboratory-confirmed cases [8]. Phylogenetic analysis of the NiV strain causing this outbreak revealed a 96.15% similarity to the Bangladesh strain [9]. Since the first report from India in 2001, the state of Kerala witnessed Nipah outbreaks in 2018 and 2021, the last one in 2024 being the sixth outbreak of the disease in the state [10, 11]. Sequential outbreaks of the disease in the Indian subcontinent have raised concerns of a re-emerging trend of this highly transmissible virus posing a major public health threat. It is one among the WHO priority pathogens that require urgent research attention for effective containment measures.

The high fatality rate (around 90% in recent outbreaks) [12] as well as the lack of effective antiviral treatment and preventive vaccines impose any studies on infectious NiV strictly restricted to BSL-4 bio-containment facilities. The absence of easily employable surveillance systems poses hindrance to prevalence studies in both humans and animals, and also in the effective and early detection of subclinical cases that may help in prevention of disease outbreaks. To circumvent this difficulty and facilitate research, several surrogate systems employing virus-like particles and pseudovirions that support studies in a BSL-2 facility have been developed.

The 18.2 kb negative-sense RNA genome of NiV codes six structural proteins and three non-structural proteins [13, 14]. The structural proteins G and F are the key envelope glycoproteins facilitating NiV binding to its cellular receptor. The glycoprotein G binds to the Ephrin B2 or B3 receptor triggering a conformational change in the fusion protein F, supporting the viral envelope fusion with the host cell membrane [15, 16]. Earlier studies with surrogate viral systems involved the use of NiV-like particles (VLPs) and pseudotyped viruses using lentivirus or Vesicular Stomatitis virus (VSV)-based systems [17–19]. The NiV VLPs that co-express the surface glycoproteins G, F, and the matrix protein M have been used in neutralization assays [3]. A pseudovirus production system for henipavirus using the MuLV packaging cell line expressing the LACZ reporter gene has been reported [13]. Exploiting the broad host range and robust replication properties of VSV, a recombinant VSV (rVSV-G*ΔG)-based system with glycoprotein G coding region deleted has been used to generate pseudotypes of several heterologous viruses including NiV [20–23]. Antibody neutralization assay using the pseudotyped NiV showed a good correlation with the gold standard PRNT assay that uses infectious NiV indicating their functional resemblance [24]. In addition to the replication defective VLP and pseudovirus-based systems, a recombinant, replication competent NiV F and G chimeric Cedar virus platform has also been generated recently as a surrogate neutralization system [25].

During their generation, all these surrogate systems need multiple transfections of plasmids encoding NiV F/G proteins or matrix protein M into the producer cells. There is a critical need for optimization of plasmid concentrations used in the transfection in order to avoid inconsistency in the surface expression of these proteins in the producer cells. This has remained challenging [26]. As a consequence, there can be variable incorporation rates of these proteins in the VLPs or pseudovirions generated and batch-to-batch variability in the infectious virus titre. There have been no major attempts made to address this issue and generate systems that can produce high titer NiV VLPs or pseudovirions. In most of the currently available surrogate systems, the reporters used as indicators of infection are fluorescence or luminescence-based, making them less amenable to high throughput automation for large scale use in surveillance or screening studies. In the present study, we address these gaps by reproducibly generating high titer VSV-based NiV pseudoviruses from F and G stably-transfected and clonally selected producer cells. A secreted alkaline phosphatase (SEAP) was incorporated as a reporter for easy readout. We also show the use of these pseudovirions in neutralization assays employing commercial antibodies as well as convalescent serum from Nipah recovered individuals.

## Materials and Methods

### Generation of NiV F and G expression vectors

Complete coding region sequences of NiV F (1640 bp; position 6650-8290) and G (1808 bp; position 8939-10747) genes derived from the National Center for Biotechnology Information (NCBI) database (GenBank Accession No.MH523642; NiV strain from the outbreak in Kerala in 2018 [9]) were used to synthesize gene constructs (BR Biochem, India). Plasmid vector backbones with a CMV promoter, pLXSN and pcDNA3.1 Hygro, were derived from the mammalian expression plasmids pLXSN-Axl (Addgene # 65222) and pHMC122 (a kind gift from Dr. Erica Ollman Saphire, La Jolla Institute of Immunology, USA). NiV F coding region was cloned to the *EcoRI*-*BamHI* linearized pLXSN vector backbone and the NiV G coding region was cloned to the *KpnI*-*HindIII* linearized pCDN3.1 Hygro vector backbone using standard cloning procedures. Chemically competent *Escherichia coli (E. coli)* DH5α strains were transformed with the ligated products and the plasmid DNA isolated from NiV F and G positive clones were used for transfection of mammalian cells.

### Cell lines

BHK-21 baby hamster kidney cells (ATCC; CCL-10) and HEK293 human embryonic kidney cells (ATCC; CRL-1573), were originally purchased from the American Type Culture Collection (ATCC) and were cultured in high glucose DMEM (Gibco).The culture media were supplemented with 10% FBS (Gibco) and 1X antimycotic antibiotic solution (Sigma) and cells were maintained at 37°C in a 5% CO2 atmosphere.

### Plasmid transfections and generation of NiV F & G stable HEK293 producer cells

HEK293 cells (3×10^5^) used for the production of pseudovirions were seeded in 6-well plates for 18h and co-transfected with the protein expression plasmids pLXSN-NiV F and pcDNA-NiVG (2µg/ml each) using polyethyleneimine (PEI; Sigma) (8µg/ml). The selection of transfected cells stably expressing both proteins was carried out by growing the cells in the presence of geneticin (625 µg/ml; for the pLXSN-NiVF) and hygromycin (50µg/ml; for the pcDNA-NiVG) for 60 days.

### Clonal selection of NiV F and G co-expressing HEK293 producer cells

HEK293 producer cells stably expressing NiV F and NiV G proteins were subjected to limiting dilution to separate single cells in individual wells of a 96-well plate to allow clonal expansion. After two rounds of selection, individual colonies were expanded to large-scale cultures and cryopreserved. Three, well-growing clones were selected (293 FG-5F6, 293 FG-7C5, and 293 FG-8C7) for further characterization and pseudovirion production.

### Flow cytometry analysis of membrane expression of F and G proteins in monoclonal HEK293 producer cells

293 FG-5F6, 293 FG-7C5, and 293 FG-8C7 cells were trypsinized from T25 flasks and 1x10^6^ cells were stained with anti-NiV F (12B2; Absolute antibody # Ab02792-1.1) and anti-NiV G (48D3; Absolute antibody # Ab2865-23.0) antibodies in flow cytometry staining buffer (2% FBS in PBS) for 1h in ice. After incubation, the cells were centrifuged at 1500rpm for 5 minutes followed by washing the pellet in FACS washing buffer (0.5% BSA in PBS) to remove the unbound antibodies. The cell pellet was then treated with mouse and rabbit secondary antibodies to anti-F and anti-G antibodies conjugated with Alexafluor 488 and AlexaFluor 597 respectively for 30 minutes in ice. Following washing, the cell pellet was resuspended in FACS staining buffer and the percentage of incorporation of F and G proteins was analyzed in a flow cytometer (BD FACS Aria Fusion).

### Immunofluorescence detection of F & G proteins in monoclonal HEK293 producer cells

3×10^4^ cells from expanded 293FG-5F6 monoclonal cells were cultured in 24-well plates overnight, washed with 1X phosphate-buffered saline (PBS), and fixed with paraformaldehyde or pre-chilled acetone-methanol (1:1) at -20⁰C for 5 min. The fixative was removed by washing the cells twice with 1X PBS and the cells were blocked using 5% BSA in PBS, with 0.05% Tween-20 for 30 min at room temperature. Subsequently, the cells were stained with anti-NiV F (12B2; Absolute antibody # Ab02792-1.1) and anti-NiV G (48D3; Absolute antibody # Ab2865-23.0) primary antibodies in blocking buffer (1:500 dilution) overnight at 4°C. Followed by washing twice with 1X PBS, the cells were incubated with Alexafluor 488 conjugated anti-mouse (CST # 4408S) and AlexaFluor 597 conjugated anti-rabbit secondary antibodies (CST, Cat # 8889S) respectively at 1:1000 dilution. DAPI (1µg/ml) was used as the nuclear stain in all experiments.

### Pseudovirus production from HEK293 producer cells

HEK293 cells (polyclonal or the 5F6 monoclonal stable cells) were seeded at a density of 6 × 10^6^ cells and cultured in T150 culture flasks for 24h. Cells were infected with rVSV G*ΔG-SEAP stocks (2×10^6^ TCID50/ ml) in OptiMEM medium at an MOI 1 for 2h at 37°C. After 2h of infection, the inoculum was removed, and the cells were washed with PBS at least 3 to 4 times to ensure the removal of trace quantities of any VSV-ΔG-SEAP virus. The cells were cultured in complete culture medium for 18–24h before the pseudovirus harvest. On the day of harvest, the culture supernatants containing the pseudotyped virions were collected, centrifuged at 3000 rpm for 20 min and stored at -80⁰C until use.

2.9 Pseudovirus infection and titration

BHK21 target cells were infected with the pseudovirions produced from either polyclonal or monoclonal HEK293 producer cells by incubating the cells with virus stocks for 2h at 37°C. 24h post-infection, cell culture supernatant was incubated at 65°C for 15 min to inactivate the endogenous alkaline phosphatase activity; and was analyzed for SEAP activity by mixing 50µl of the supernatant with 150 µl of pNPP substrate solution (TCI, Cat. # N1109). After incubation for 10 min, absorbance at 405nm was measured using Promega GloMax® Discover Microplate Reader.

For virus titration, the 50% tissue culture infectious dose (TCID50) was determined by the Reed and Muench method [27] for each batch of the pseudovirions. Single-use aliquots were used to avoid inconsistencies in titer resulting from repeated freeze-thawing. 100 µl of serial two-fold dilutions of the pseudovirus aliquots were used to infect the target BHK-21 cells for 2h at 37°C in 96-well plates as described above. Infectivity was analyzed by SEAP assay. The cell supernatant harvested from un-transfected HEK293 cells, infected with rVSV-G*ΔG-SEAP virus served as a negative control in the SEAP assay. As described [26] a positivity cut-off for pseudovirus infection was defined as an OD405 value greater than 3SD (standard deviation) of the OD405 value of cell controls, and used in the calculation of the TCID50.

### Detection of F & G protein incorporation in NiV pseudovirions by immunofluorescence

For immunofluorescence analysis, 3 ×10^4^ BHK-21 target cells were cultured in 24-well plates and the cell monolayer was infected with 1000 TCID50 N-PV(FG)-5F6-SEAP pseudovirions for 30min at 25°C. BHK-21 cells, infected with rVSV G*ΔG-SEAP virus, were used as controls. The infected cells were washed twice with PBS and fixed using acetone:methanol (1:1) at -20⁰C followed by immunostaining to microscopically visualize NiV F and G proteins using specific primary and secondary antibodies as described above.

### NiV Pseudovirus-based Neutralization assays

BHK-21 target cells (2×10^4^) were seeded in 96 well plates in DMEM containing 10 % FBS and 1X antimycotic antibiotic solution and incubated at 37 °C in a CO2 incubator. 250 TCID50 N-PV(FG)-5F6-SEAP pseudovirions were mixed with serial dilutions of commercial NiV F and G antibodies or convalescent human sera from Nipah recovered patients [28] and incubated at 25°C for 1h. Rabishield (17C7), a NiV non-neutralizing antibody or serum from two healthy individuals was used as the negative control in the neutralization assays. Serum samples were heat inactivated at 56°C for 30 min before using in the assays to inactivate the complement. The pseudovirus-sera / antibody mixture was added to the monolayer of BHK-21 cells. After 2h infection at 37 °C, the mixture was removed from the cells, replenished with DMEM containing 2% FBS and 1X antimycotic antibiotic solution, and incubation was continued for 24h. The cell culture supernatants were harvested; and incubated at 65°C for 15 min to inactivate the endogenous alkaline phosphatase activity and the secreted alkaline phosphatase (SEAP) was measured as described above.

### Statistical analysis

Statistical analyses were performed using Graphpad prism 8.0 software. Pearson’s correlation analysis was used to quantify the pixel-to-pixel proportionality in the signal levels of the two channels in immunofluorescence experiments. A Student’s t-test was used to compare the significance between two groups. P value <0.05 was considered statistically significant.

## Results

The general scheme of experiments followed to generate and characterize the NiV pseudovirions is shown in Fig.1.

**Fig. 1.**
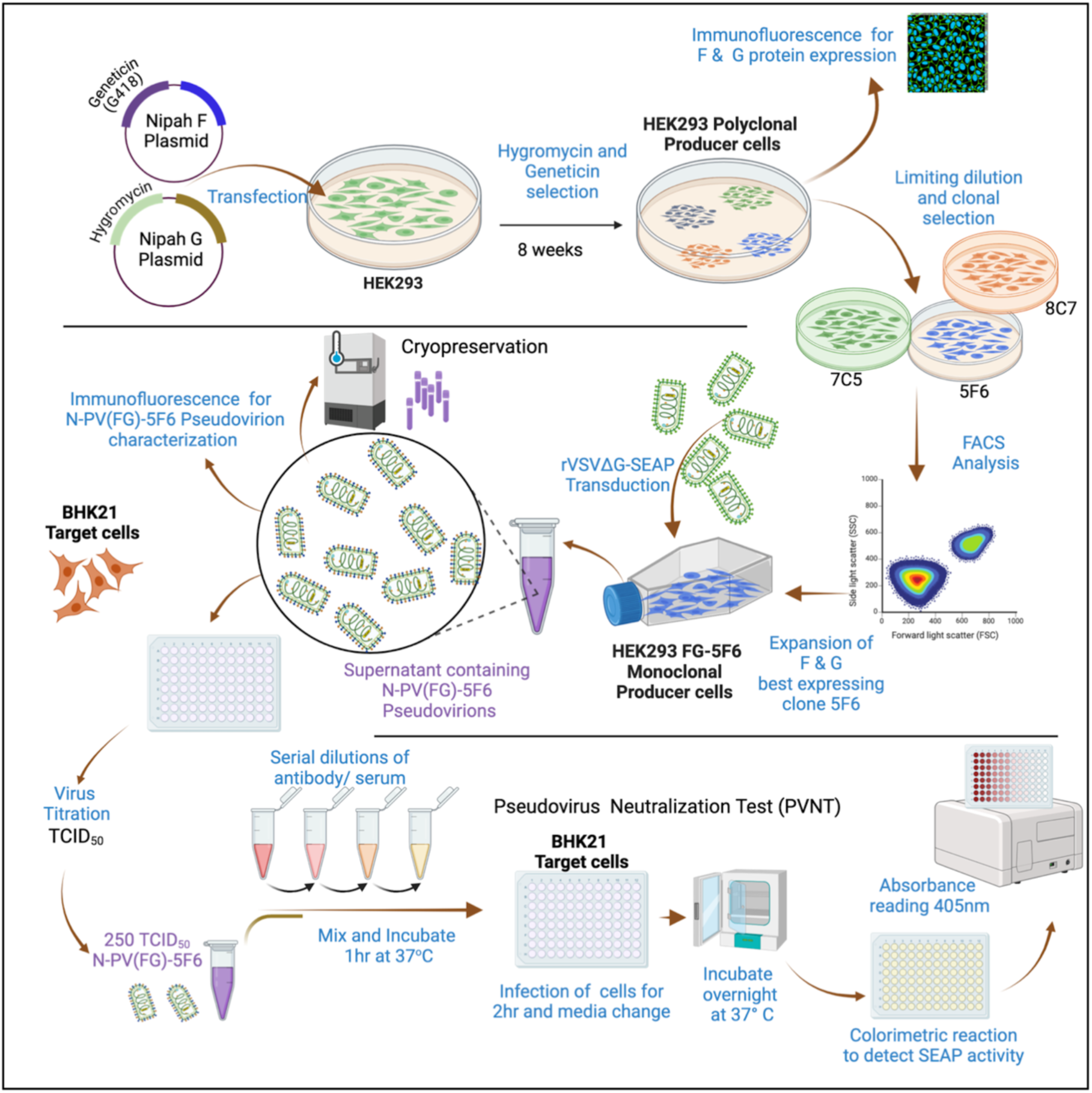
Scheme of experiments for generation and characterization of NiV pseudovirions.

### Generation of pseudovirions from polyclonal HEK293 producer cells stably expressing NiV F and G protein

HEK293 polyclonal producer cells were co-transfected with the pLXSN-NiVF and pcDNA3.1-NiVG clone plasmids and selected with appropriate antibiotics to generate stable cells. The expression of F and G proteins in the transfected cells was confirmed by immunofluorescence analysis using specific antibodies (Fig.2A). Infection of these cells with 2×10^6^ TCID50/ ml rVSV-G*ΔG-SEAP virus stock led to the generation of N-PV(FG)-SEAP pseudovirions 24h post-transfection. This was evidenced by enhanced SEAP activity in the supernatants of BHK21 target cells infected with this virus compared to mock infection control (Fig.2B).

**Fig. 2.**
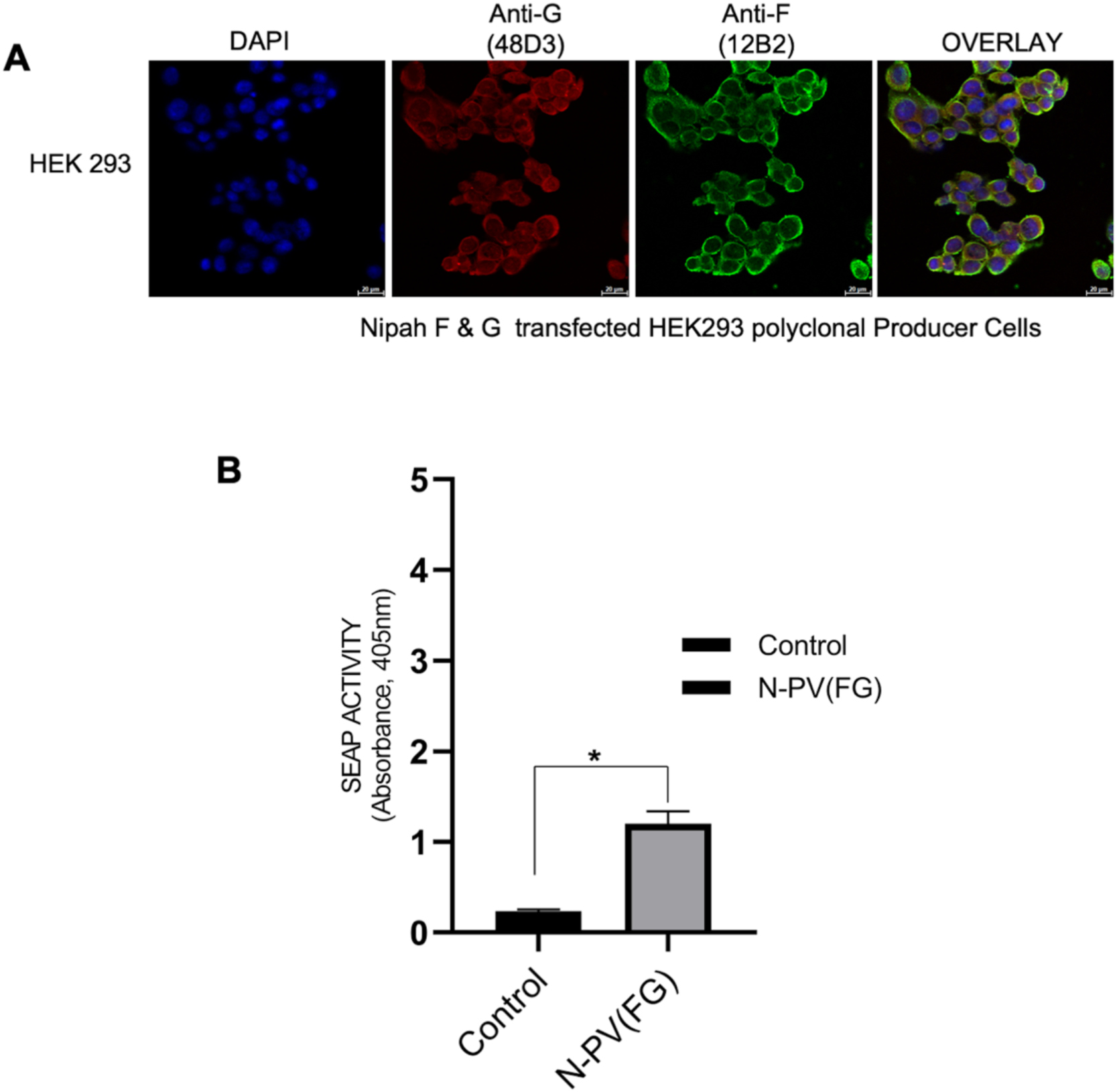
Generation of stably transfected HEK293 polyclonal producer cells and characterization of the infectivity of the N-PV (FG) pseudovirions generated. **A.** Immunofluorescence analysis of HEK 293 cells stably expressing NiV G and F proteins stained with primary antibodies to Nipah G (48D3) and Nipah F (12B2) respectively (Magnification, 60X; Scale bar, 20µm). **B.** The SEAP activity measurements of N-PV(FG) compared with the un-transfected infected HEK 293 control. Data presented as Mean+/- SD; *, P<0.05. Unpaired Student t-test was used for statistical analysis comparing the two groups. Experiment was done in duplicates and data presented as Mean+/-SD of three independent experiments (N=6).

### Clonal selection and characterization of NiV F and G co-expressing monoclonal HEK293 producer cells

Stably co-transfected HEK293 cells with NiVF and G expressing plasmids were subjected to successive rounds of selection under the selection pressure of hygromycin and geneticin. These cells were subjected to limiting dilution for the selection of monoclones of producer cells. Three well defined and rapidly growing monoclones selected from 96-well plates were expanded in 24-well plates and subjected to FACS analysis. The percentage of incorporation of NiV F and G proteins in the three monoclones 293FG-5F6, 293FG-7C5, and 293FG-8C7 was found to be 90.5%, 88.9%, and 29.6 % respectively (Fig. 3A). The uniform expression levels and membrane colocalization of the NiV F and G proteins in 293FG-5F6 monoclonal cells was also evident immunofluorescence analysis (Fig. 3B). Hence, this clone was further expanded and selected as producer cells for large-scale pseudovirus production.

**Fig. 3.**
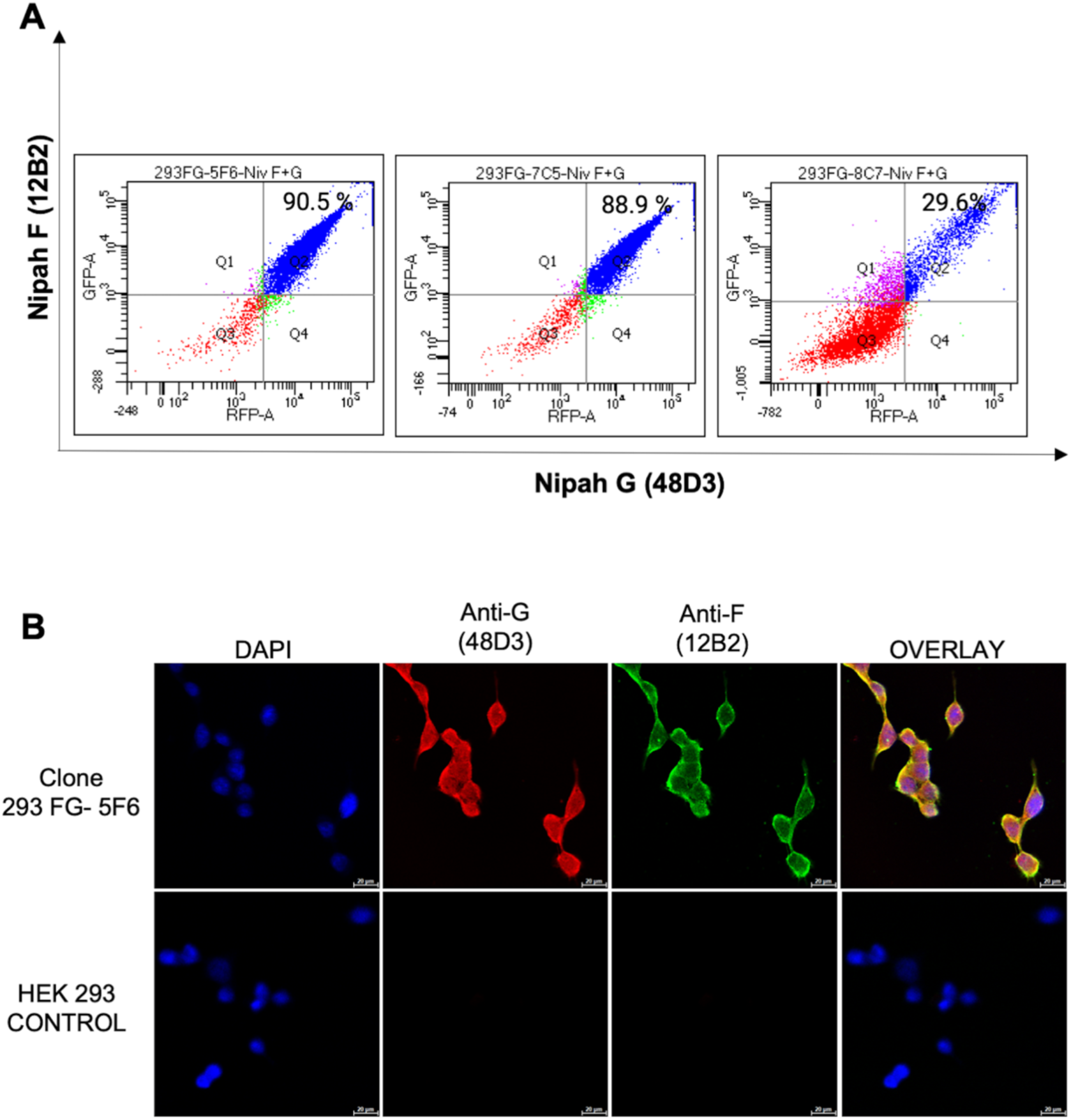
Analysis of co-expression of NiV surface glycoproteins F and G in clonally expanded stable HEK293 cell lines. **A.** NiV F and G protein expression in clonal cells. Suspensions of cells from expanded monoclones 293FG-5F6, 293FG-7C5 and 293FG-8C7 were stained with NiV anti-G and anti-F antibodies as described in the methods; and analysed by FACS. The percentage incorporation of G and F proteins was found to be 90.5%, 88.9 %and 29.6%, respectively. **B**. Immunofluorescence analysis of NiV G and F protein co-expression in 293FG-5F6 cells. The fixed cells were stained with primary antibodies 48D3 and 12B2 and secondary antibodies Alexafluor 597 and Alexafluor 488 respectively for G and F proteins and the membrane expression of the proteins were viewed under a confocal microscope. Untransfected HEK293 cells served as mock control in A and B. DAPI was used as the nuclear stain (Magnification, 60X; Scale bar, 20µm).

### Reproducible generation of high-titre NiV pseudotyped virions from 293FG-5F6 cells

We next investigated whether the 293FG-5F6 monoclonal producer cells can generate high titer N-PV(FG)-5F6 pseudovirions when infected with rVSV-G*ΔG-SEAP stock virus.

Significantly high levels of SEAP activity was observed in target BHK-21 cell culture supernatants when infected with N-PV(FG)-5F6, compared to the infection with N-PV(FG) pseudovirions produced from polyclonal producer cells (Fig. 4A). This indicated a high titer pseudovirion production by the 293FG-5F6 cells.

**Fig. 4.**
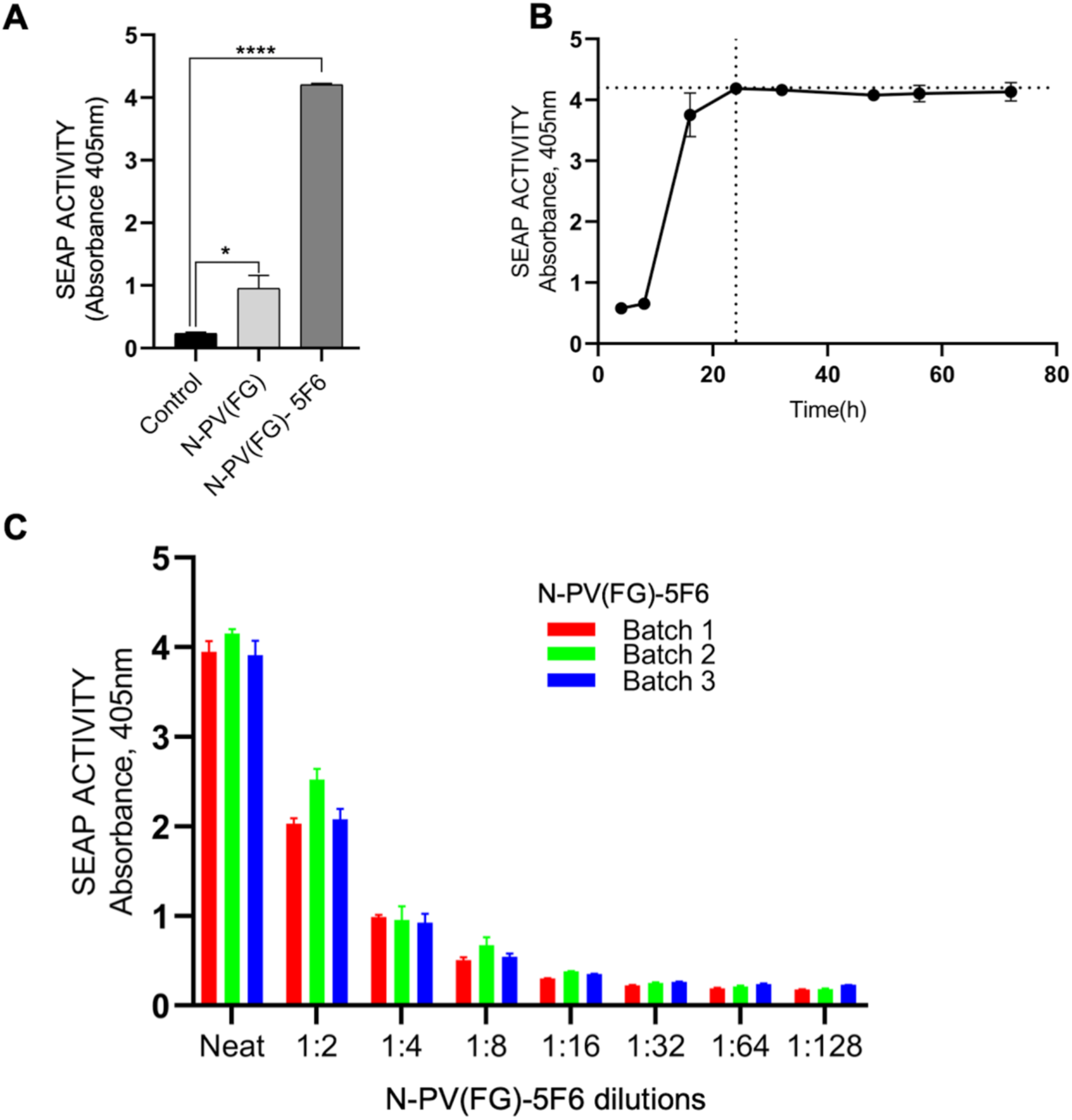
Characterization of N-PV(FG)–5F6 pseudovirions. **A.** Comparison of SEAP activity in culture supernatants from infected BHK-21 target cells. Cells were infected with identically produced pseudovirion harvests N-PV(FG) and N-PV(FG)-5F6 respectively from polyclonal and monoclonal HEK293 producer cells. Supernatant from un-transfected HEK293 cells infected with rVSV (G*△G)SEAP was used for infecting control cells. The values are the Mean±SD of duplicate values from three experiments *, P<0.05; **** P<0.0001; Unpaired Student t-test was used for statistical analysis comparing the two groups. **B.** Kinetics of SEAP activity in BHK-21 target cell supernatnats post-N-PV(FG)-5F6 infection. Data points indicate the time points 4,8, 16, 24, 32, 48, 56 and 72h of collection post-infection. The 24h time-point fixed for subsequent experiments is indicated by the dotted line. Each value is a Mean±SD of triplicate values from two experiments **C.** Validation of reproducibility of N-PV(FG)-5F6 pseudovirion production under identical conditions. Three batches of pseudovirions harvested from HEK293(FG)-5F6 monoclonal producer cells were used to infect BHK-21 target cells in various dilutions. SEAP activity in the culture supernatant was measured at 405nm and plotted. Each value indicates Mean±SD of triplicate values from three independent experiments (N=9).

Kinetic analysis of the SEAP activity in culture supernatants from N-PV(FG)-5F6 pseudovirions infected BHK-21 target cells revealed that the activity reaches maximum by 24h post-infection and thereafter remains stable (Fig.4B). So this time-point was chosen as the end-point in subsequent biological assays. In order to check the reproducibility of pseudovirion production, three batches of N-PV(FG)-5F6-SEAP viruses were generated from 293FG-5F6 clonal cells under identical conditions of rVSV-G*ΔG-SEAP infection. These three stock viruses had similar infectivity to the target BHK-21 cells as indicated by the SEAP activity in the culture supernatants from infections done at different dilutions (Fig.4C).

### Detection of NiV F and G proteins on N-PV(FG)-5F6 pseudovirions by immunolocalization with specific antibodies

To identify whether NiV F and NiV G protein co-expressed on N-PV(FG)-5F6 pseudovirions, immunolocalization was carried out on BHK-21 target cells post infection with specific antibodies and was analyzed by confocal microscopy. Superimposition of the green and red fluorescence indicated the co-distribution of F and G proteins respectively in the pseudovirions produced. This was well apparent in the yellow color of the overlay image of N-PV(FG)-5F6 pseudovirion infected cells (Fig.5A). However, mock infection with rVSV-G*ΔG-SEAP virus gave no signal indicating the specificity of the detection (Fig.5A). Also, there was significantly low expression of the G protein on the N-PV(FG) pseudovirions. Scatter plot analysis of the fluorescence expression on pseudovirions revealed distribution of the points around a central straight line passing through the origin in the case of N-PV(FG)-5F6, as against a skewed distribution on N-PV (FG), indicating a uniform expression of these proteins in the former (Fig.5B&C). Quantification of the mean fluorescence intensity expression levels of F & G proteins reconfirmed the equal levels of incorporation of both the proteins in N-PV(FG)-5F6 pseudovirions, compared to that in N-PV(FG) (Fig.5 D&E). F and G protein expression levels in N-PV(FG)-5F6 pseudovirions had a Pearson’s coefficient of 0.93 whereas in N-PV(FG), it was only 0.83 in correlation analysis indicating a significantly better uniform incorporation in the former.

**Fig. 5.**
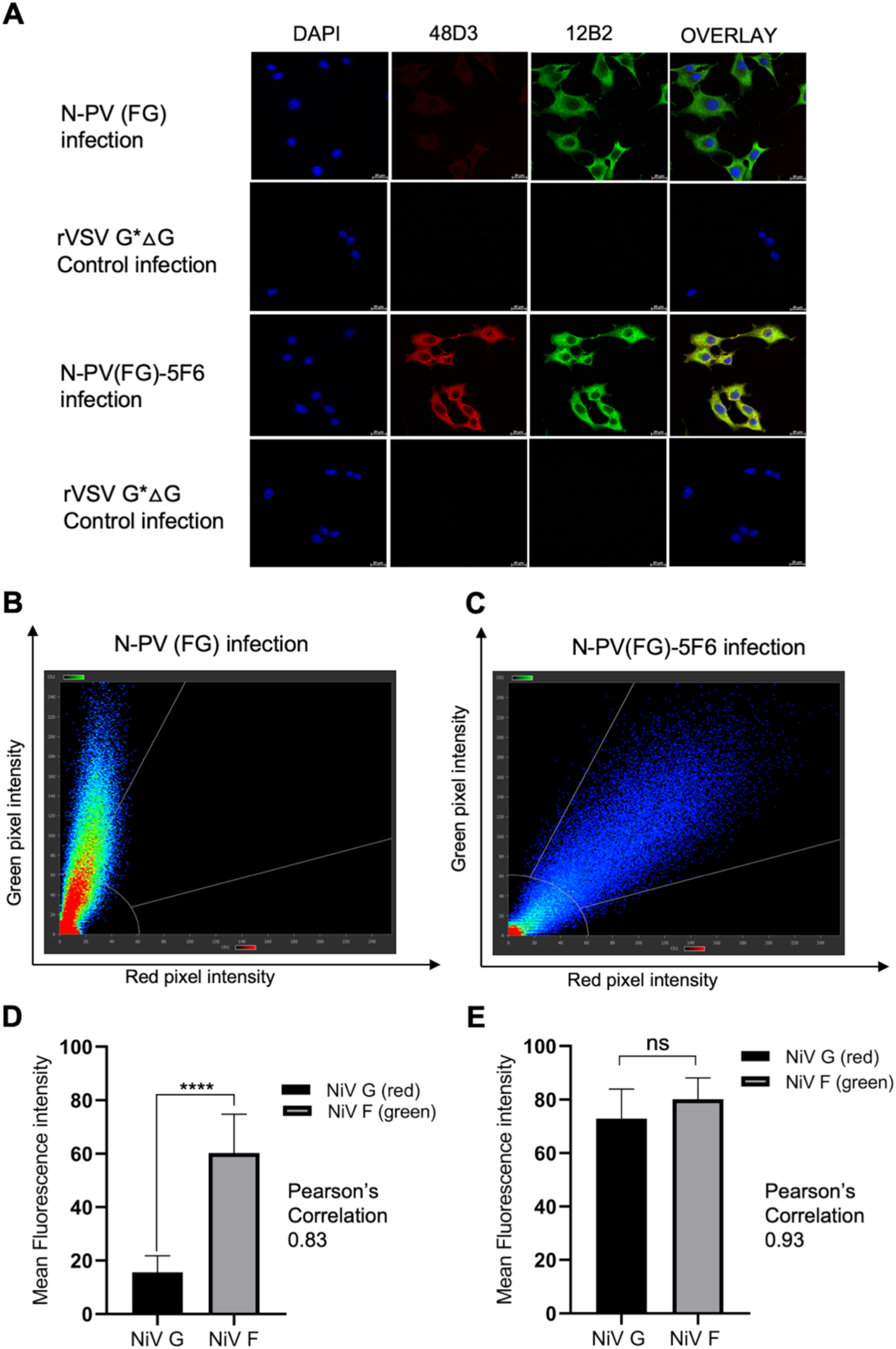
Evaluation of uniform expression of F &G proteins on NiV Pseudovirions. **A.** Localization of N-PV(FG)-5F6 pseudovirions on target cells post-infection by immunostaining with anti-NiV F & G antibodies and confocal microscopy. BHK-21 cells were infected with pseudovirions for 30 min at 25°C to facilitate virus attachment. Cells were fixed in acetone: methanol (1:1) and stained with primary antibodies 48D3 and 12B2; and secondary antibodies Alexafluor 597 and Alexafluor 488, respectively to visualize the G and F proteins on the attached pseudovirions. **B & C.** Scatterplot of green and red pixel intensities respectively corresponding to NiV F and G staining on N-PV(FG) and N-PV(FG)-5F6 pseudovirions. The points of the scatterplot distributed more towards the green fluorescence probe in N-PV(FG) compared to a proportional co-distribution in N-PV(FG)-5F6 infection where the points of scatterplot cluster around a central straight line**. D& E.** Quantification of co-localization using Pearson’s correlation coefficient (PCC). A PCC value of 0.83 in N-PV(FG) showed that the probes did not overlap in a fixed proportion whereas a high PCC value of 0.93 for N-PV(FG)-5F6 probes reflected proportional co-distribution. The values are the Mean±SD of 14 region-of-interests (ROIs) fileds from two independent experiments. **** P<0.0001; ns-Not significant; Unpaired Student t-test was used for statistical analysis comparing the two groups.

### Pseudovirus neutralization assays using commercial anti-NiV antibodies and convalescent serum from Nipah infection recovered individuals

The usefulness of the N-PV(FG)-5F6 pseudovirions in functional assays were confirmed by neutralization assays. Commercial anti-NiV F and G antibodies gave 100 % neutralization of the pseudovirions (Fig.6A). The neutralization titer decreased with decreasing antibody concentration, confirming a dose-dependent neutralizing activity and its specificity. The 17C7 non-neutralizing antibody used as negative control did not neutralize the N-PV(FG)-5F6 pseudovirions, further confirming the specificity of the observations.

**Fig. 6.**
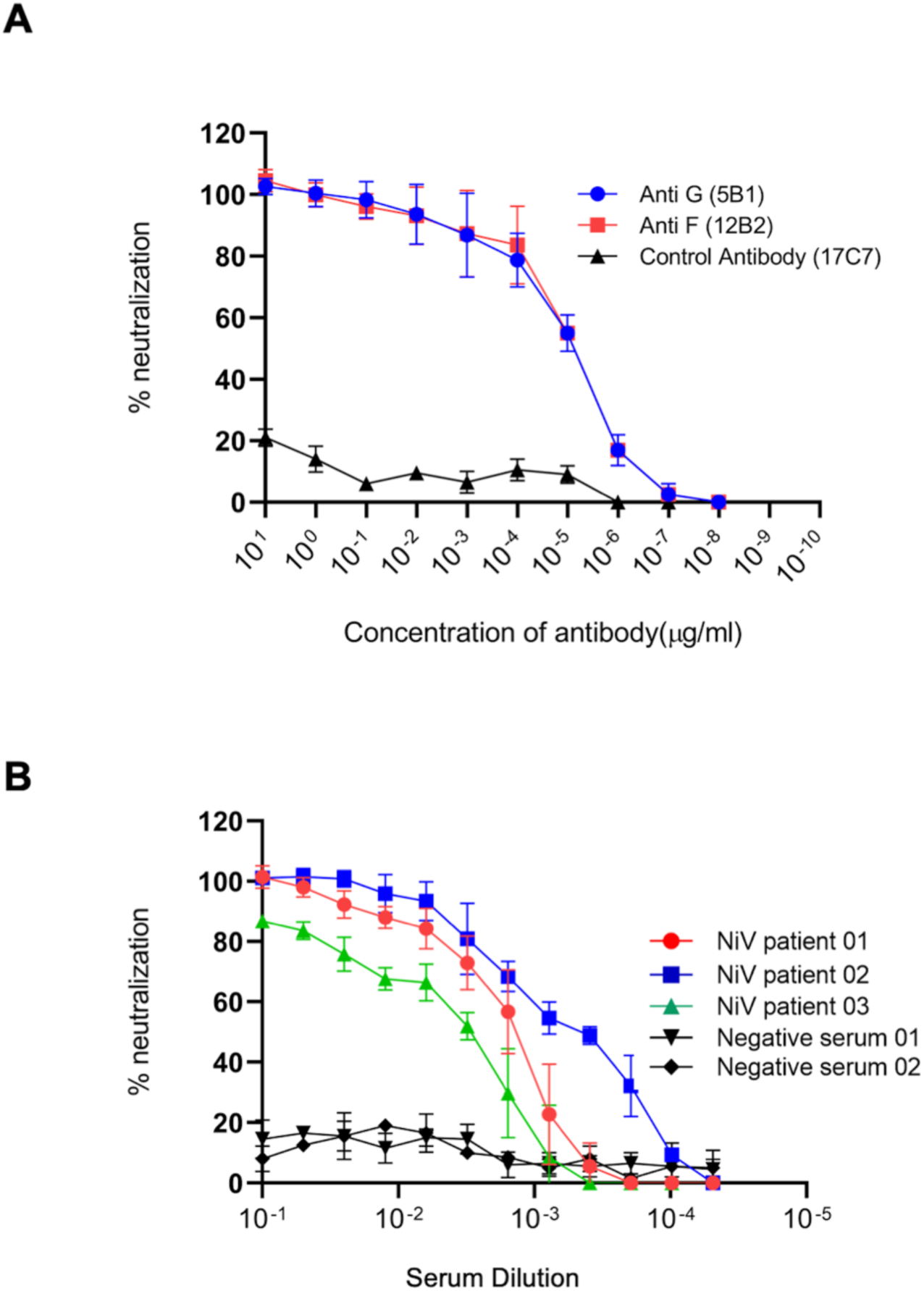
N-PV(FG)-5F6 pseudovirion neutralization by commercial anti-Nipah antibodies and convalescent serum from Nipah recovered patients. **A**. Neutralization curves of N-PV(FG)-5F6 pseudovirions neutralizing commercial Anti-G and Anti-F antibodies. 250 TCID_50_ pseudotyped virions were pre-incubated with increasing concentrations of neutralizing antibodies to Nipah F and G structural proteins. A non-neutralizing antibody mAb 17C7 (Rabishield) was used a negative control in the experiments. Experiment was done in duplicates and each data point presented is a Mean+/- SD of three independent experiments (N=6). **B**. Assay of NiV positive patient sera using N-PV(FG)-5F6 pseudovirions. Neutralization curves of N-PV(FG)-5F6 pseudovirions neutralizing NiV positive sera named as NiV patient 01, NiV serum 02 and NiV serum 03. 250 TCID_50_ pseudotyped virions were pre-incubated with increasing dilutions (1:10, 1:20, 1:40, 1:80, 1:160, 1:320, 1:640, 1:1280, 1:2560, 1:5120, 1:10,240, and 1:20,480) of patient sera. Experiment was done in triplicates and each data point is a Mean+/- SD of three independent experiments (N=9).

The susceptibility of the N-PV(FG)-5F6 pseudovirions to neutralization by human antibodies were evaluated using convalescent serum samples from three Nipah recovered subjects (collected during the Nipah outbreak that occurred in the Kozhikode district of Kerala, India in the year 2023;). Serum samples (NiV patient 01, 02, and 03) were diluted to 1:10 followed by 2-fold serial dilutions; and used in the assays. All the three samples showed neutralizing activity against NiV pseudovirions. While NiV serum 01 and 02 could completely neutralize the pseudoviruses, NiV03 gave more than 80% neutralization at a serum dilution of 1:10 (Fig. 6B). The percentage of neutralization showed a declining trend with increasing serum dilutions. Sera from two normal individuals used as controls in the assay did not neutralize the N-PV(FG)-5F6 pseudovirions (<20 %). From these results, it could be concluded that the established high throughput system is sensitive and specific and could be used for serosurveillance in Nipah endemic regions.

## Discussion

NiV spill over and human disease outbreaks are sporadic and unpredictable. The lack of commercial rapid diagnostic or serological tests prevent early detection and large-scale epidemiological investigations. Like for other BSL-4 pathogens such as Ebola virus (EBOV), surrogate systems, such as virus-like particles and pseudoviruses, have been developed for NiV. However, unlike EBOV and similar viruses that have only one surface glycoprotein (GP) that facilitate infection and to be included in these systems [29], NiV has two surface glycoproteins F and G that are to be incorporated. In all the previous reports on NiV VLP-based or pseudovirus-based systems, the authors attempted transient co-transfections of F and G expressing plasmids with multiple optimization attempts of outer membrane and backbone plasmid concentration ratios to obtain their co-expression. This resulted in the unreliable incorporation of these proteins in the progeny particles, making the approach challenging and less reproducible [17–19, 30]. In a recent study, a cell line stably expressing the NiV F protein was infected with a recombinant VSV expressing NiV G to produce high-titre pseudovirions [31]. Though the study had confirmed expression of the F & G proteins in the transfected and rVSV infected cell lines, it did not analyse the efficiency of incorporation of both the proteins in the generated pseudovirions.

In the present study, to overcome the above difficulties, we generated producer cells that stably express the F and G proteins after transfection. Initial cloning of the F and G genes was carried out in mammalian expression vectors that have different antibiotic selection markers, G418 and hygromycin; and stable polyclonal cells were generated after multiple rounds of selection for two months. As shown in Fig.2A, immunofluorescence analysis confirmed that the NiV F and G proteins are expressed on these producer HEK293 cells. When pseudovirions generated from these cells were used to infect the target BHK-21 cells and the viral particles were visualized by immunostaining, both F and G proteins could be detected. However, when these images were overlayed and analysed, only a poor protein co-localization could be observed (Fig.5A). This indicated that the N-PV(FG) pseudoviral particles generated from these polyclonal cells were still heterogenous even after prolonged selection for stable expression of both the F and G proteins.

To further improve the pseudovirion generation and make our assay more reproducible, the stable producer cells were subjected to clonal selection by limiting dilution to ensure single cell per well, which were further expanded and analyzed. Analysis of three clones (293FG-5F6, 293FG-7C5, and 293FG-8C7) from the clonal selection demonstrated a significant variation in the rate of expression of both the proteins; and only the clone 293FG-5F6 that showed more than 90% expression of both the proteins were chosen for the establishment of Nipah pseudovirus system. These clonal cells had an almost equal levels of F and G protein membrane expression as seen in the immunostaining (Fig.3B)). Pseudovirion particles from these producer cells had a conspicuous co-localization of both the proteins (Fig.5A) and these cells showed consistency in production of pseudovirions with uniform infectivity (Fig.4C). Co-localization of the fluorescence probes of NiV F and G have been seldom quantified in any of the previous studies, thus making this assay system novel and reliable. To the best of our knowledge, this is the first attempt to make stable producer cells co-expressing the NiV F and G proteins that can be exploited for production of uniform batches of NiV pseudovirions.

Pseudovirus-based systems are finding increased use in antibody neutralization assays and antiviral screening assays for identifying virus entry inhibitors [32]; and simpler and high throughput-adaptable assays are a pre-requisite for their large-scale use. The present study used secreted alkaline phosphatase (SEAP) as a reporter in the NiV pseudovirions generated. Earlier studies have used SEAP-reporter systems for the development of pseudovirus systems for other viruses such as human papillomavirus [33, 34]. Except for one report that used SEAP [20], all other previous studies on NiV surrogate systems, including the recent study [31] used either fluorescence or luminescence-based reporter assays for monitoring infection. SEAP-based assays are non-destructive and provide about 10-fold higher sensitivity than conventional luciferase-based systems [21, 35]. It also offers high throughput adaptability as well as easy monitoring of the pseudovirion infection using conventional ELISA readers providing easier implementation in large-scale surveillance programs. We found that the SEAP activity becomes detectable by 18h post-infection and reaches a peak by 24h post-infection in NiV pseudovirion infected target cells (Fig.4B), facilitating an early readout in biological assays using this system.

As surrogate systems need not mimic the structural organization and functional properties of the surface glycoproteins of natural viruses in all aspects, it is imperative to test the functionality of the pseudovirus particles generated. Neutralization by specific antibodies is a key parameter that indicates their functionality and is a useful property that helps to assess virus-specific immunity in a population. The use of pseudoviruses for evaluating antibody-mediated neutralization has been described in previous studies [36]; and also, earlier reports have indicated a good correlation of the results from these assays with conventional PRNT assays using natural, infectious viruses [24]. However, the use of too excess pseudoviruses can give false negative results in neutralization assays and there is a need to use optimal virus concentration to obtain reliable and reproducible results. We found that a dilution of pseudovirion stock that gives a titer of approx. 250 TCID50 gave a dynamic working range OD405 of 1.00-0.2 in our PVNT assays starting with virus controls to negative/ background controls. Accordingly, this dilution of the stock N-PV(FG)-5F6 pseudovirions was used in all the neutralization assays.

Antibodies generated in a mouse system (commercial anti-NiVF and G mAbs) as well as human convalescent serum were used to evaluate the pseudovirion functionality in neutralization assays. Both antibodies showed a dose-dependent neutralization of the NiV pseudovirions while the negative control antibodies did not neutralize the virus as expected (Fig.6 A & B). The assay detect the neutralizing antibodies in 1:10 diluted serum samples indicating its comparable performance with conventional virus neutralization assays. NiV pseudovirions were designed using the sequences of the Bangladesh strain of virus that caused the Nipah outbreak in Kerala in 2018 [9]; and recent studies have pointed out that the 2023 Nipah outbreak, from where the serum samples were collected, was also caused by the same strain of NiV . There were differences in the level of neutralization of the three patient samples used pointing out that the assay detects quantitative differences in the specific antibody titers in each of these patients.

## Conclusions

The present study developed a robust and reproducible system for generating VSV-based NiV pseudovirions. These pseudovirions uniformly express F and G proteins and can be effectively used in virus neutralization assays. Further validation of the assay in a larger set of samples will make this rapid and high throughput-adaptable assay for cost-effective use in large scale epidemiological screening studies in simple laboratory settings. This assay will also be significantly useful in therapeutic monoclonal antibody and vaccine response evaluation as well as in drug discovery studies against Nipah.

## Abbreviations

NiV: Nipah Virus
VLP: Virus-like particle
PVNT: Pseudovirus Neutralization Test
VSV: Vesicular Stomatitis Virus
FACS: Fluorescence Activiated Cell Sorting
SEAP: Secreted Alkaline Phosphatase
BSL: Biosafety Level
TCID: Tissue Culture INfective Dose
BHK: Baby Hamster Kidney
HEK: Human Embryo Kidney
ATCC: American Type Culture Collection
MOI: Multiplicity of Infection

## Acknowledgments

Authors acknowledge Mr.Gopikrishnan K for the excellent technical support with confocal microscopy and flow cytometry; and Dr. Erica Ollman Saphire, La Jolla Institute of Immunology, USA for sharing reagents for the study. The authors are also grateful to the Additional Chief Secretary (Health), Government of Kerala for granting permission to obtain convalescent samples for the study.

## Author Contributions

GRV : Pseudovirion generation, clonal selection, virus neutralization assays, confocal and flow cytometry experiments; and writing the first draft of the manuscript.

VV : Pseudovirion assays, ultracentrifugation experiments.

SS : Generation of the NiV F & G clones.

SSL : Cell culture experiments and virus titration

PP : Convalescent serum sample collection

SA : Convalescent serum sample collection

APM : Convalescent and healthy serum sample collection

NKP : Convalescent serum sample collection

ES : Conceived the study, carried out overall supervision, arranged funding and edited and finalised the manuscript. All authors approved the final version of the manuscript.

## Data Availability

All data generated or analysed during this study are included in this published article.

## Declarations

### Ethics approval and consent to participate

The study protocol was approved by the Institutional Human Ethics Committees (IHECs) of the Government Medical College, Kozhikkode, Kerala and the Institute of Advanced Virology, Thiruvananthapuram, Kerala (Approval No.IHEC/IAV/2023/02 dated October 27, 2023). Human serum samples were collected from Nipah recovered as well as healthy individuals, strictly adhering to the protocols approved by the IHEC. Informed, written consents were obtained from all the participants.

